## Appendices for "Genome concentration limits cell growth and modulates proteome composition in *Escherichia coli*"

#### Appendix 1

#### Estimation of the average genome concentrations

**Cell area, cell volume and growth rate**. The cell width $w$, cell length $L$, and cell area $A$ under different conditions were measured in our experiments. From these values, we estimated the cell volume by $V(L,w) = \left(\frac{4\pi}{3}\right)\left(\frac{w}{2}\right)^3 + \pi\left(\frac{w}{2}\right)^2(L-w)$, assuming that the *E. coli* cell poles are hemi-spheres. For different culture conditions, the normalized growth rate $\lambda = \frac{1}{A}\frac{dA}{dt}$ was estimated by tracking the cell area increase during time-lapse imaging, using the UNet segmentation algorithm (Zhou et al., 2020) with customized MATLAB scripts (see https://github.com/JacobsWagnerLab). The doubling time $\tau_{DB}$ is calculated by $\tau_{DB} = log2/\lambda$. All values are listed in Appendix 1 – Table 1 below.

**DNA concentration**. To estimate the average DNA concentration for cells under balanced growth, we assumed that the cell volume grows exponentially during the cell cycle and that the DNA content follows a specific DNA replication pattern (see Appendix 1- figure 1A). We used published information on the pattern of a fluorescent fusion to the DNA replication marker SeqA (Govers et al., 2024) to extract information about DNA replication. If a cell cycle event (i.e., initiation of DNA replication) happened at the division cycle percentile $\theta$ (from 0 to 100% corresponding to cell birth and division, respectively), this value can be inferred from population statistics with a suitable cell cycle marker. A demograph (Hocking et al., 2012) can be constructed by sorting cells by length, allowing one to identify the demograph percentile $\phi$ (from 0 to 100%) of the cell cycle event (see Appendix 1-figure 1B). To adjust for the non-uniform age distribution in exponentially growing populations, a conversion formula between cell cycle percentile and demograph percentile was used, namely, $\theta = 1 - \frac{\log(2-\phi)}{\log(2)}$ (Wold et al., 1994).

To take into account the overlapping DNA replication rounds in cells growing in rich medium (M9glyCAAT), we defined $\theta_1$ and $\theta_2$ as the cell cycle percentiles for initiation and termination of DNA replication, respectively. These values were estimated from the fluorescent SeqA pattern on the demograph (see Appendix 1-figure 1B), where the corresponding demograph percentiles were estimated to be $\phi_1 = 50\%$ and $\phi_2 = 60\%$. Using the conversion formula, we obtained $\theta_1 = 41.5\%$ and $\theta_2 = 51.5\%$. In the case of non-overlapping DNA replication in nutrient-poorer media (M9gly and M9ala), we define $\theta_B$ and $\theta_{B+C}$ as the cell cycle percentiles for initiation and termination of DNA replication, respectively. These values were estimated from the fluorescent SeqA pattern in the demograph (see Appendix 1-figure 1B), and the corresponding demograph percentiles were estimated. For the M9gly condition, we have $\phi_B = 0\%$ and $\phi_{B+C} = 64\%$ and this corresponded to $\theta_B = 0\%$ and $\theta_{B+C} = 56\%$. For the M9ala condition, we have $\phi_B = 41\%$ and $\phi_{B+C} = 77\%$ and this corresponded to $\theta_B = 33\%$ and $\theta_{B+C} = 71\%$.

Using these values, we extracted the DNA content $Z(t)$ and cell volume $V(t)$ during the division cycle (see Appendix 1 – figure 1B) and calculated the mean DNA concentration $[Z]_{avg}$ by averaging $Z(t)/V(t)$ across the division cycle.

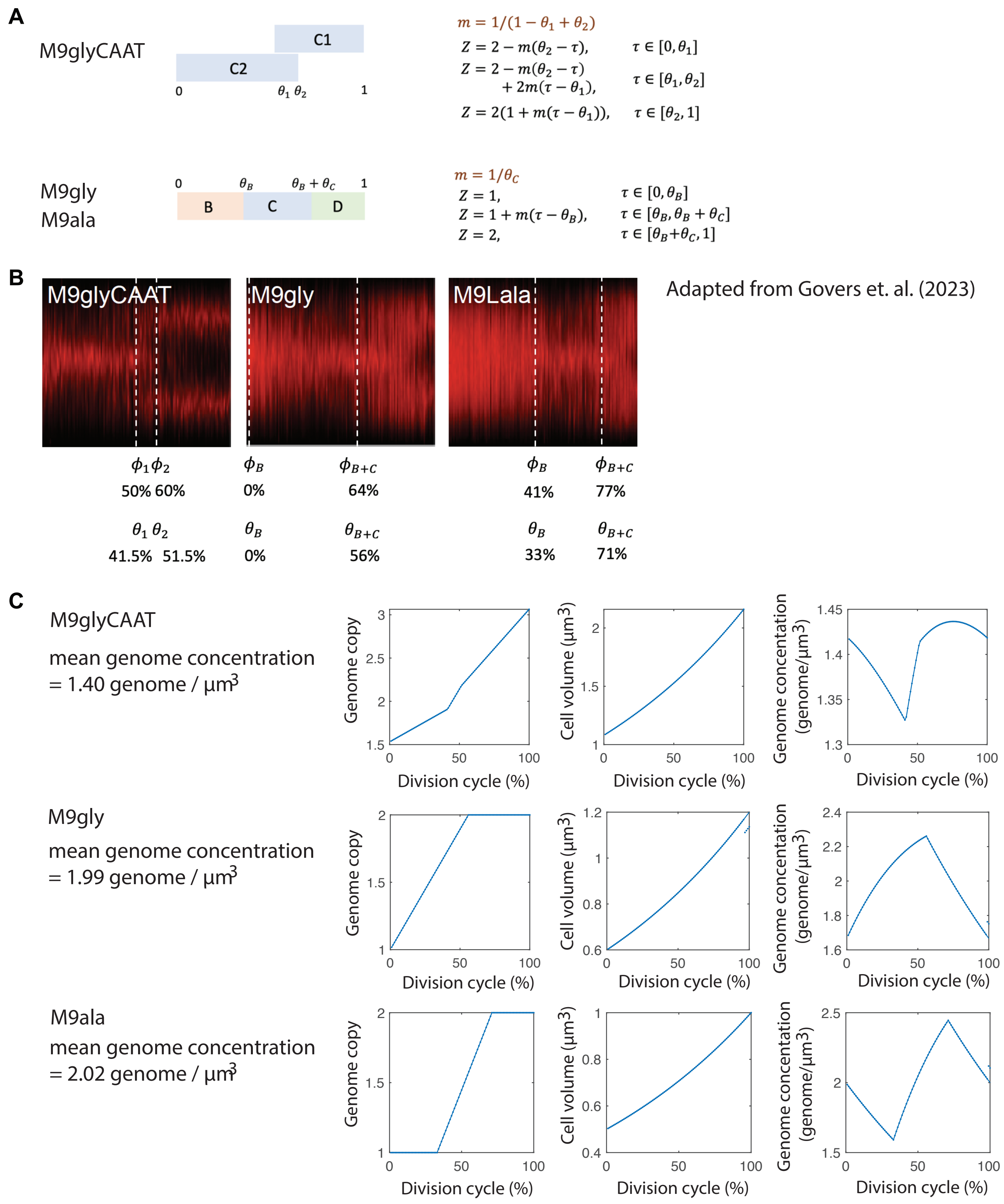

**Appendix 1 – Figure 1. Estimation of DNA concentration. (A)** DNA replication pattern for different medium conditions. **(B)** Extraction of the genome content, cell volume and genome concentration along the division cycle. **(C)** Genome copy, cell volume, and genome concentration along the division cycle.

**Appendix 1 – Table 1. Sizes, DNA concentrations, and growth rates of cells in different growth media.**

| Symbol | Parameter | Growth condition | | | Source |
|---|---|---|---|---|---|
| | | M9glyCAAT | M9gly | M9ala | |
| $A$ | Mean cell area (µm$^2$) | 2.48 | 1.51 | 1.40 | Measured in this work |
| $w$ | Mean cell width (µm) | 0.80 | 0.69 | 0.62 | Measured in this work |
| $L$ | Mean cell length (µm) | 3.25 | 2.44 | 2.51 | Measured in this work |
| $V$ | Mean cell volume (µm$^3$) | 1.50 | 0.83 | 0.70 | Estimated from cell length and width |
| $V_{ini}$ | Cell volume at birth (µm$^3$) | 1.08 | 0.60 | 0.50 | $V_{ini} = V/(2log2)$ for exponential-growing population |
| $\lambda$ | Normalized growth rate (1/min) | 0.0173 | 0.0069 | 0.0039 | Measured in this work |
| $\tau_{DB}$ | Doubling time (min) | 40 | 100 | 175 | Estimated from $\lambda$ |
| $[Z]_{avg}$ | Average DNA concentration under balanced growth (genome/µm$^3$) | 1.40 | 1.99 | 2.02 | Calculated from cell cycle averaging of the DNA content and cell volume. |
| $[Z]_{ini}$ | DNA concentration of 1N cells at birth (genome/µm$^3$) | 0.92 | 1.67 | 2.00 | Calculated from $V_{ini}$ with the assumption of initial DNA content = 1 genome. |
| $c$ | Cell volume:protein ratio ($10^6$ µm$^3$) | 0.283 | 0.339 | 0.530 | See Appendix 1 - Tables 1 and 2 for $V$ and $Y$ estimates, respectively. |
| $c'$ | Cell area:protein ratio ($10^6$ µm$^2$) | 0.469 | 0.616 | 1.061 | See Appendix 1 - Table 1 for $V$ and $A$ estimates. |

#### Appendix 2

#### Estimation of the model parameters

To simulate our ODE models (see Eq. 1 and Eq. 2 in the main text), we needed to estimate the kinetic constants during transcription and translation ($r_1, r_2, K_1, K_2, \delta$, see Table 1 in the main text for details) and the ratio constants $c = \frac{\text{cell volume}}{\text{protein number}}$, $c' = \frac{\text{cell area}}{\text{protein number}}$. We also needed the values of protein and mRNA molecules in newborn cells ($X_{ini}, Y_{ini}$) as initial conditions for the simulations, as well as the DNA concentration of exponentially growing wild type cells, $[Z]_{avg}$. For 1N cell, the DNA content was fixed as 1 copy per cell. In this section, we explain how these parameters were curated from the literature.

**Kinetic constants for transcription and translation.** Our experiments were conducted in M9glyCAAT at 37°C. To estimate physiological parameters (mRNA and protein concentrations, etc.), we used datasets from the literature and interpolated using the doubling time (or growth rate) of our experimental condition as a reference. Three datasets (Balakrishnan et al., 2022; Bremer et al., 2003; Bremer and Dennis, 2008) were used for estimating the following kinetic constants:

<u>(i) (Bremer and Dennis, 2008):</u> The authors curated physiological values of mRNA, protein, RNAP and ribosome numbers in exponentially growing cells with doubling times $\tau = 20, 24, 30, 40, 60, \text{and } 100\ min$ (all growing at 37°C using different carbon sources). In our experiments under M9glyCAAT, *E. coli* cells have a doubling time $\tau = 40\ \ min$. Hence, we used the values of $\tau = 40\ min$ from Bremmer and Dennis (2008) for our simulations.

<u>(ii) (Bremer et al., 2003):</u> The authors calculated the fractions of RNAPs associated with rRNA (including *rrnP1* and *rrnP2*), mRNA (including constitutive and repressible genes), or under paused condition. These numbers were obtained for M9Glucose with casamino acid (with normalized growth rate 0.0277 $min^{-1}$) and M9Gly (with normalized growth rate 0.0115 $min^{-1}$), both at 37°C. To estimate the fraction of RNAPs engaged in mRNA synthesis, we used the formula is shown in Appendix 2 - Figure 1G, which we solved for the normalized growth rate of our growth condition.

<u>(iii) (Balakrishnan et al., 2022):</u> The authors measured the mRNA degradation rates for normal and carbon-limited conditions, and showed that the mRNA degradation rate is not sensitive to growth rate. Therefore, we used the same mRNA degradation rate to model all our medium conditions (see Appendix 1 - Table 4).

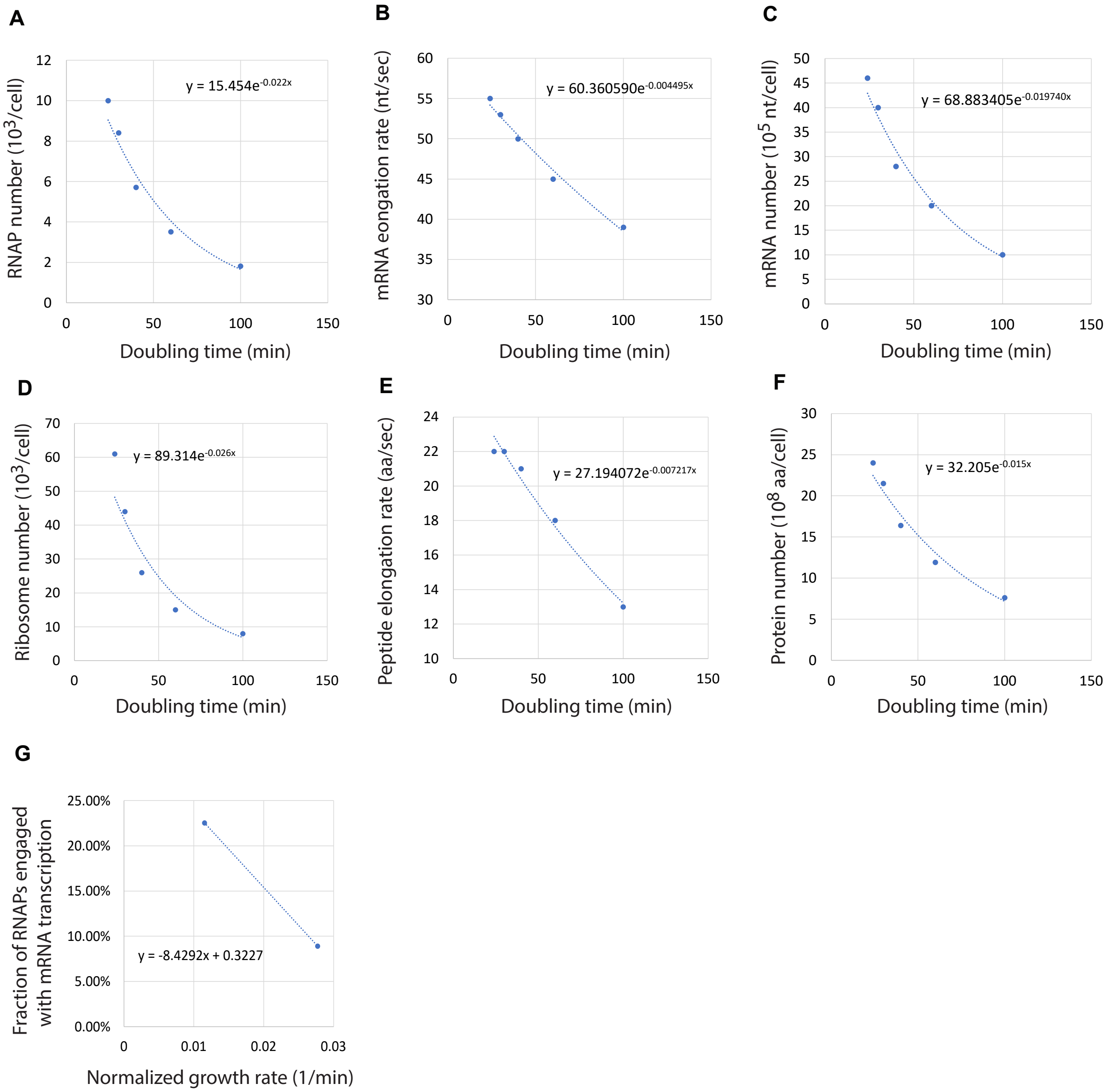


**Appendix 2 – Figure 1. Interpolation and extrapolation of parameters.** The parameters of the ODE models reflecting different medium conditions were calculated using the fitted formulae. The obtained values are summarized in Appendix 2 – Table 1.

**Appendix 2 – Table 1. mRNA and protein numbers per cells and bulk rates of transcription and translation.** This table describes how kinetic constants were estimated from the literature. Parameters $X_{ini}, Y_{ini}, r_1, r_2$ were used in ODE simulations. We assumed exponential growth and used the relation $M_{ini} = M/(2log2)$ where $M_{ini}$ is the biomass of a newborn cell and $M$ is the average cellular biomass in the population (Koch and Schaechter, 1962). The bulk transcription rate $r_1$ was defined as $r_1 = \frac{\text{mRNA synthesis rate}}{\text{\# of proteins in the cell}}$, and the bulk translation rate $r_2$ was defined as $r_2 = \frac{\text{protein synthesis rate}}{\text{\# of proteins in the cell}}$. Values were estimated from a previous study (Bremer and Dennis, 2008). For our estimations, we assumed that the average protein length is 310 amino acids and that the average mRNA length is about 1 kb (Ishihama et al., 2008).

| Symbol | Parameter | Value | Source |
|---|---|---|---|
| $X$ | Number of mRNAs (per cell) | 2800 | Estimated from (Bremer and Dennis, 2008) |
| $Y$ | Number of proteins ($10^6$/cell) | 5.29 | Estimated from (Bremer and Dennis, 2008) |
| $X_{ini}$ | Number of mRNAs for a newborn cell | 2020 | Calculated as $X_{ini} = X/(2log2)$ |
| $Y_{ini}$ | Number of proteins for a newborn cell ($10^6$) | 3.82 | Calculated as $Y_{ini} = Y/(2log2)$ |
| $n_{RNAP}$ | Number of RNAPs in the cell ($10^3$) | 5.7 | Estimated from (Bremer and Dennis, 2008) |
| $n_{protein}$ | Number of proteins in the cell ($10^6$) | 5.29 | Estimated from (Bremer and Dennis, 2008) |
| $\theta_{RNAP}$ | Fraction of RNAPs relative to the total number of proteins | 0.00108 | $\theta_{RNAP} = n_{RNAP}/n_{protein}$ |
| $\beta_{mRNA}$ | Fraction of RNAPs involved in mRNA synthesis | 0.18 | Estimated from (Bremer et al., 2003). |
| $T_{RNAP}$ | Mean transcription time to generate one mRNA molecule (min) | 0.33 | Estimated from (Bremer and Dennis, 2008) |
| $r_1$ | Bulk transcription rate ($10^{-3}$/min) | 1.99 | $r_1 = \theta_{RNAP}\beta_{mRNA} /T_{RNAP}$ |
| $n_{ribo}$ | Number of ribosomes in a cell ($10^3$) | 26 | Estimated from (Bremer and Dennis, 2008) |
| $n_{protein}$ | Number of proteins in a cell ($10^6$) | 5.29 | Estimated from (Bremer and Dennis, 2008) |
| $\theta_{ribo}$ | Ratio between ribosome number and protein number ($10^{-3}$) | 4.91 | $\theta_{ribo} = n_{ribo}/n_{protein}$ |
| $T_{ribo}$ | Mean translation time for one protein molecule (min) | 0.25 | Estimated from (Bremer and Dennis, 2008) |
| $r_2$ | Bulk translation rate ($10^{-3}$/min) | 20.0 | $r_2 = \theta_{ribo} /T_{ribo}$ |

**Appendix 2 – Table 2. DNA and mRNA affinity constants ($K_1$, $K_2$).** The active fraction of RNAPs and ribosomes, $\alpha_{RNAP}$ and $\alpha_{ribo}$, are given by the formulae $\alpha_{RNAP} = \frac{[Z]}{K_1+[Z]}$ and $\alpha_{ribo} = \frac{[X]}{K_2+[X]}$ where $[Z]$ and $[X]$ are the DNA concentration and the mRNA concentration in the cells, respectively. To infer the parameters $K_1$, $K_2$, we used the values of $\alpha_{RNAP}$, $\alpha_{ribo}$, $[Z]$ and $[X]$ of wild-type cells determined in our study and back calculated the values of $K_1$, $K_2$.

| Symbol | Parameter | Value | Source |
|---|---|---|---|
| $\alpha_{RNAP}$ | Active fraction of RNAPs | 0.5 | Measured in this work |
| $[Z]$ | DNA concentration (genome/ µm$^3$) | 1.40 | Estimated in Appendix 1 – Table 1 |
| $K_1$ | DNA affinity of RNAPs (genome/ µm$^3$) | 1.40 | Calculated as $K_1 = \frac{1-\alpha_{RNAP}}{\alpha_{RNAP}}[Z]$ |
| $\alpha_{ribo}$ | Active fraction of ribosomes | 0.75 | Measured in this work |
| $[X]$ | mRNA concentration (1/ µm$^3$) | 1867 | $[X] = X/V$ |
| $K_2$ | mRNA affinity of ribosomes (mRNA/µm$^3$) | 622 | Calculated as $K_2 = \frac{1-\alpha_{ribo}}{\alpha_{ribo}}[X]$ |

**Appendix 2 – Table 3. Estimation of the mRNA degradation rate.** The mRNA degradation data were obtained from an experimental study (Balakrishnan et al., 2022).

| Symbol | Parameter | Value | Source |
|---|---|---|---|
| $\delta$ | mRNA degradation rate (1/min) | 0.964 | (Balakrishnan et al., 2022) |
| $\tau_d$ | mRNA lifetime (min) | 1.04 | Calculated by $\tau_d = 1/\delta$ |
| $\tau_H$ | mRNA half life (min) | 0.72 | Calculated by $\tau_H = (\log 2)/\delta$ |

### Appendix 3

#### Model B, an alternative ODE model that includes the experimentally observed change in RNAP concentration in 1N cells

For the first ODE model A, we assumed that the active fraction of RNAPs follows Michaelis-Menten type kinetics as $\alpha_{RNAP} = \frac{[Z]}{K_1+[Z]}$. This model A only considered DNA concentration [Z] but ignored the concentration of RNAP. In 1N cells, we experimentally found that the RNAP concentration increases with cell area (Figure 3A). An increasing RNAP concentration could potentially decrease $\alpha_{RNAP}$ due to an increased competition between RNAP molecules for DNA-binding sites (i.e., promoters). To take into account the increase in RNAP concentration in 1N cells, we considered a more sophisticated kinetic model B. In this model, RNAP (denoted by $P$) is classified into three states:

1. Promoter-bound state ($P_p$)
2. Elongation state ($P_e$)
3. Free state ($P_f$)

The promoter (denoted by $Q$) is classified into two states:

1. RNAP-bounded state ($Q_b$)
2. Free state ($Q_f$)

The transition between these states is illustrated in the diagram below.

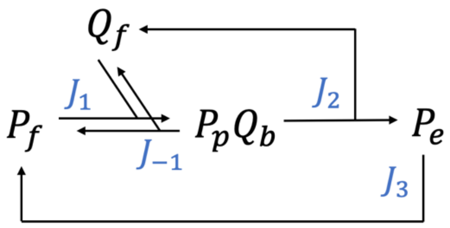


The fluxes between these states are:

$$J_1 = k_1[P_f][Q_f]$$
$$J_{-1} = k_{-1}[P_p] = k_{-1}[Q_b]$$
$$J_2 = k_2[P_p] = k_2[Q_b]$$
$$J_3 = k_3[P_e]$$

Here, $k_1$ and $k_{-1}$ are the ON and OFF constants between free RNAPs and unbounded promoters, respectively. $k_2$ is the rate constant for transcription initiation, while $k_3$ is the rate constant that is inversely proportional to the transcription time. Under steady state, we have

$$0 = \frac{d[P_f]}{dt} = (-J_1) + J_{-1} + J_3$$

$$0 = \frac{d[P_p]}{dt} = J_1 - J_{-1} - J_2$$

$$0 = \frac{d[P_e]}{dt} = J_2 - J_3$$

Solving the equations and let $[P] \equiv [P_f] + [P_p] + [P_e]$ and $[Q] \equiv [Q_f] + [Q_b]$, we get

$$[P] = [P_f]\left(1 + \frac{k_1}{k_{-1}+k_2}[Q_f] + \frac{k_1k_2}{(k_{-1}+k_2)k_3}[Q_f]\right) \qquad [4\text{-}1]$$
$$[Q] = [Q_f]\left(1 + \frac{k_1}{k_{-1}+k_2}[P_f]\right) \qquad [4\text{-}2]$$

We assume a separation of time scale such that $[P]$ (concentration of all RNAPs) and $[Q]$ (concentration of all promoters) are constant. Solving [4-1], [4-2], we obtain

$$[P_f] = \sqrt{C_1^2 + C_2} - C_1$$

with $C_1 \equiv \frac{1+A[Q]-B[P]}{2B}$, $C_2 \equiv \frac{[P]}{B}$, $A \equiv \frac{k_1}{k_{-1}+k_2}\left(1+\frac{k_2}{k_3}\right)$, $B \equiv \frac{k_1}{k_{-1}+k_2}$.

Note that the active fraction of RNAPs is $\alpha_{RNAP} \equiv 1-\frac{[P_f]}{[P]}$, and the promoter occupancy is $\theta_{occ} \equiv \frac{[Q_b]}{[Q]}$. Therefore, the active fraction of RNAPs can be expressed as function of $[P]$ and $[Q]$, i.e.,

$$\alpha_{RNAP}([P],[Q]) = 1-\left(\sqrt{f_1^2+f_2}-f_1\right)$$

with $f_1 \equiv \frac{1+A[Q]-B[P]}{2B[P]}$, $f_2 \equiv \frac{1}{B[P]}$, $A \equiv \frac{k_1}{k_{-1}+k_2}\left(1+\frac{k_2}{k_3}\right)$, $B \equiv \frac{k_1}{k_{-1}+k_2}$.

**(a) Estimation of [P], [Q] and rate constants for model B**

We needed to estimate the concentrations of RNAPs and promoters for cells in the M9glyCAA condition. From the constant $c = 0.283\ (\mu m^3/protein)$, the proteome fraction of RNAP $\theta_{RNAP} = 1.08 * 10^{-3}$, and the average cell volume $V = 1.5\ \mu m^3$ (Appendix 1 – Tables 1 and Appendix 2 – Table 1), we obtain $[P] = 2.12 \times 10^3 \left(\frac{\#}{\mu m^3}\right) = 3.39\ \mu M$.

The number of promoters in the cells was assumed to be $n_Q = 3000$ promoter (Wang et al., 2013). In M9glyCAA condition, the average genome copy is $[Z]_{avg} = 1.4 \left(\frac{genome}{\mu m^3}\right)$ (Appendix 1 – Table 1). This corresponds to $[Q] = 4.2 \times 10^3 \left(\frac{\#}{\mu m^3}\right) = 6.72\ \mu M$.

Under physiological condition, the concentrations of RNAPs and promoters are in the micromolar range. We obtained the following kinetic constants from the literature (Bettridge et al., 2023): $k_{-1} = 1.1\ s^{-1}$, $k_2 = 0.012\ s^{-1}$, $k_3 = 0.0083\ s^{-1}$. To estimate the remaining parameter $k_1$, we note that the wild-type cells in M9glyCAAT condition has $\alpha_{RNAP} = 0.5$ (Appendix 2 – Table 2) and this value corresponds to $k_1 = 7.5 \times 10^4\ M^{-1}s^{-1}$ in model B.

**(b) Estimation of the cell size-dependence of RNAP concentration**

From our experimental measurements, the RNAP concentration [P] increases linearly with the cell area of 1N cells (orange line in Figure 3A). This can be described by the following phenomenological equation:

$$[P] = [P_0] + b(A - A_0)$$

If we let $1\ AU' = 10^5 AU$, then the slope $b$ is estimated to be 0.0786 ($AU'/\mu m^2$).

To develop a biophysical model for this dependency, we rationalized that 1N cells can sense the dilution of DNA and upregulate their proteome fraction of RNAP ($\theta^*_{RNAP}$) accordingly. This suggests that $\theta^*_{RNAP}$ is a function of DNA concentration $[Z]$.

Since in wild-type condition $\theta^*_{RNAP}$ is only about 0.1% of the proteome (Bremer and Dennis, 2008), we assumed that the ratio between cell volume and total protein number ($c$) is a constant and that the ratio between cell area and the total protein number ($c'$) is also a constant. To apply these assumptions to 1N cells, we note that

$$[P] = \frac{\#RNAP}{V} = \frac{\#RNAP}{cY} = \frac{1}{c}\theta^*_{RNAP}$$

$$[P_0] = \frac{\#RNAP_0}{V_0} = \frac{\#RNAP_0}{cY_0} = \frac{1}{c}\theta_{RNAP}$$

where $\#RNAP_0$ represents the number of RNAPs in an average sized wild-type cell, and $[P_0], V_0, A_0, Y_0$ represent the RNAP concentration, the cell volume, the cell area and the protein number, respectively. For 1N cells, the genome copy per cell is always 1, hence

$$A = \frac{c'}{c}V = \frac{c'}{c}\frac{1}{[Z]}$$

$$A_0 = \frac{c'}{c}V_0 = \frac{c'}{c}\frac{1}{[Z_0]}$$

Here, $[Z_0]$ is the average DNA concentration of 1N cells under normal cell size $V_0$ (before DNA dilution). For reference, we have $\theta_{RNAP} = 1.08 \times 10^{-3}$, $V_0 = 1.5\ \mu m^3, A_0 = 2.5\ \mu m^2$, $Z_0 = 0.67\ \mu m^{-3}$, $c = 0.28 \times 10^6\ \mu m^3$, $c' = 0.47 \times 10^6\ \mu m^2$ (see Appendix 1 – Table 1 and Appendix 2 – Table 2). Substituting the above relation by the linear phenomenological equation $[P] = [P_0] + b(A - A_0)$, we have

$$\theta^*_{RNAP} = \theta_{RNAP} + bc(A - A_0) = \theta_{RNAP} + bc'\left(\frac{1}{[Z]} - \frac{1}{[Z_0]}\right)$$

To estimate the coefficient $b$ from experimental measurements, we needed to convert the RNAP concentration from AU' to ($\#/\mu m^3$). We noticed that the RNAP concentration in wild-type cells is $\theta_{RNAP}c^{-1} = 4.15\ \times 10^3$ ($\#/\mu m^3$). This corresponds to 0.75 (AU') in the experimental measurements. Therefore, we have 1 (AU') = $5.53 \times 10^3$ ($\#/\mu m^3$). From the experimental measurements, $b = 0.0786$ ($AU'/\mu m^3$), hence for 1N cells, we have $b = 430$ ($\frac{\#}{\mu m^5}$). For multi-N cells, we simply assumed that $b = 0$ since based on our experimental measurements (Figure 3A, blue line), $[P]$ does not depend on cell area.

**(c) Comparison between fraction of active RNAPs predicted by models A and B**

| Model A | Model B |
|---|---|
| $\frac{dX}{dt} = r_1 \alpha_{RNAP} Y$<br><br>$r_1 = \theta_{RNAP} \beta_{mRNA} / T_{RNAP}$<br><br>$\alpha_{RNAP} = \frac{[Z]}{K_1 + [Z]}$ | $\frac{dX}{dt} = r_1^* \alpha_{RNAP} Y$<br><br>$r_1^* = [\theta_{RNAP} + bcc'(Y - Y_o)]\, \beta_{mRNA} / T_{RNAP}$<br><br>$\alpha_{RNAP}([P],[Q]) = 1 - \left(\sqrt{f_1^2 + f_2} - f_1\right)$<br><br>$f_1 \equiv \frac{1+A[Q]-B[P]}{2B[P]},\ \ f_2 \equiv \frac{1}{B[P]}$<br><br>$A \equiv \frac{k_1}{k_{-1}+k_2}\left(1 + \frac{k_2}{k_3}\right),\ \ B \equiv \frac{k_1}{k_{-1}+k_2}$<br><br>$[P] = \frac{1}{c}[\theta_{RNAP} + bcc'(Y - Y_o)]$<br>$[Q] = n_Q [Z]$ |

To compare the two models, we first calculated the $\alpha_{RNAP}$ for both cases. Under physiological (WT-like cell sizes) conditions both the concentration of both RNAPs [P] and promoters [Q] are in micromolar range. In Appendix 3- figure 1, we see that both models give comparable landscape of $\alpha_{RNAP}$. Note that model A has no dependence on [P], while model B shows some negative dependence on [P].

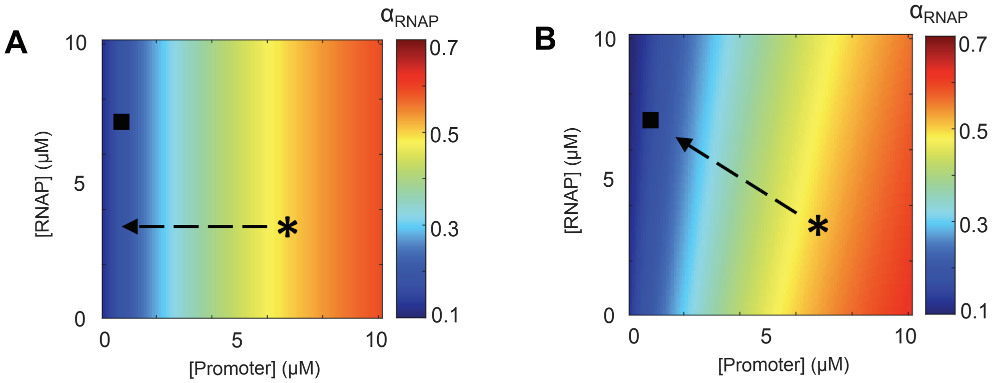


**Appendix 3 – Figure 1. Comparing changes in active RNAP fraction between ODE models A and B.** The two-dimensional colormaps show the values of active RNAP fraction ($\alpha_{RNAP}$) under different promoter and RNAP concentrations. (**A**) Using the formula of model A with M9glyCAAT parameters. (**B**) Using the formula of model B with M9glyCAAT parameters. The asterisk and filled square symbols indicate the promoter and RNAP concentration of wild-type cells and 1N cells (with 10 $\mu m^2$ cell size), respectively. The dashed arrow indicates the trajectory of promoter and RNAP concentration under DNA-limitation growth in the model.

#### Appendix 4

#### Fitting experimental data with the optimized ODE models

**Parameter optimization for the ODE model.** In Appendix 2, we curated the parameters from the literature and interpolated to match our experimental conditions. Based on the experimental methods used, the measured values may sometimes differ by 50% or more. In addition, we used interpolation and extrapolation to extract some values. Therefore, it is conceivable that our estimated values could deviate from the real physiological values. Since the goal of our mathematical model is to capture the general growth behavior of DNA limitation, we allowed the parameters to vary within a realistic range (see below) to improve the simultaneous fitting to six experimental datasets (growth rate, fraction of active RNAPs, and fraction of active ribosomes for both multi-N/WT and 1N cells).

We developed a scheme to obtain optimized parameters that fit the experimental data (as shown in Figure 5A-C and Figure 5 – figure supplement 1A-C). For both model A and B, the optimization scheme focused on six parameters: $r_1$ (bulk mRNA synthesis rate), $r_2$ (bulk ribosome synthesis rate), $K_1$ (RNAP affinity to DNA), $K_2$ (ribosome affinity to mRNAs), $\delta$ (mRNA degradation rate), $c$ (ratio between cell volume and protein number). For the other parameters such as DNA concentration $[Z]$ and the number of protein and mRNA molecules for newborn cells ($X_{ini}$, $Y_{ini}$), we fixed the values curated from the literature.

For the first-round optimization, we used the curated parameters from the literature (denoted as initial parameter set, see Table 2 in the main text) and allowed parameters ($r_1, r_2, K_1, K_2, \delta, c$) to randomly vary by $\pm 50\%$. By repeating the same procedure 30,000 times, we explored the neighborhood of the initial parameter space. For each randomly varied parameter set, we simulated the ODE and compared the simulated results to our six experimental datasets. We calculated the error for each parameter set (see next section "objective function for error minimization") and chose the parameter that yielded the smallest error. This chosen parameter set was then used for the next round of optimization.

For the $i^{th}$ round ($i = 2, \ldots$) of optimization, we used the optimized parameters from the previous round and randomly varied the parameters by $\pm 5\%$. For each round, we repeated the same procedure 3,000 times and chose the parameter set that gave the smallest error. The optimization iteration was stopped when the error reduction reached a plateau (relative error reduction was less than 1%), yielding to the final optimized parameter set. For model A, the optimization terminated in 9$^{th}$, 8$^{th}$ and 21$^{st}$ rounds, respectively. For model B, the optimization terminated in the third round. Comparison between initial and optimized parameter sets are shown in Appendix 4 – Figure 1.

**Objective function for error minimization**. In this section, we explain the details for calculating the error function. For each medium condition, six datasets were included in the optimization scheme:

1. Growth rate of multi-N (or WT) cells
2. Growth rate of 1N cells
3. Fraction of active RNAPs in multi-N (or WT) cells
4. Fraction of active RNAPs in 1N cells
5. Fraction of active ribosomes in multi-N (or WT) cells
6. Fraction of active ribosomes in 1N cells

Note that we used the multi-N cell dataset for M9glyCAAT condition and the WT cell datasets for the M9Gly and M9Ala conditions. Our goal was to choose the parameter set that minimized the error between ODE simulations and experiments. For each dataset $k = 1, \dots, 6$, we defined the *mean relative error* by $E_k \equiv \sum_{j=1}^{N_k} \frac{\left|y_j^{data} - y_j^{simu}\right|}{y_j^{simu}}$, where $y_j^{data}$ is the $j^{th}$ data point in the experiment, $y_j^{simu}$ is the value predicted by the ODE model corresponded to this data point, and $N_k$ is the number of data points in the experiments. To take into account the six different datasets, we defined the objective function by $E \equiv \sum_{k=1}^{6} w_k E_k$ where $w_k$ is the weighting factors that balance the contribution between datasets. We set $w_k = 1$ for all datasets, since the multi-N and 1N datasets sets have comparable range of cell area in M9glyCAAT.

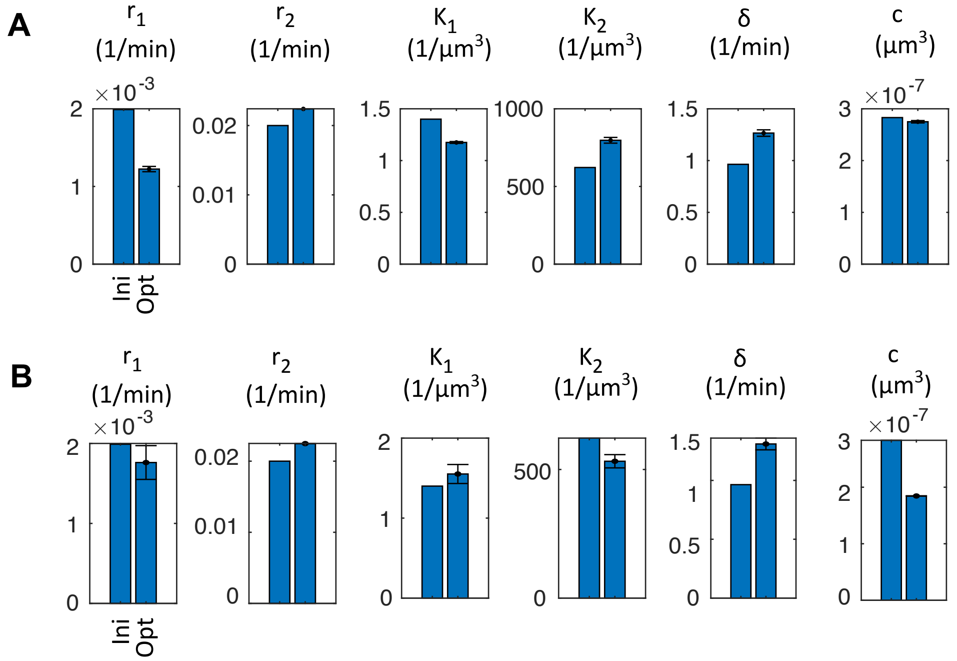


**Appendix 4 – Figure 1.** (**A**) Parameters used in model A. (**B**) Parameters used in model B.
